# A songbird karyotype: cytogenetic confirmation of a migration-associated region rich in olfactory receptor genes

**DOI:** 10.64898/2026.05.04.721007

**Authors:** Violeta Caballero-Lopez, Dmitrij Dedukh, Diana Ekman, Ondřej Kauzál, Max Lundberg, Linda Odenthal-Hesse, Estelle Proux-Wéra, Radka Reifová, Jiří Reif, Marie Altmanová, Vladimir Trifonov, Staffan Bensch

## Abstract

The field of genetics of bird migration advances, driven by exponential refinements of sequencing and tracking technologies. In willow warblers (*Phylloscopus trochilus*), a complex repeat-rich region named MARB (Migration Associated Repeat Block) has recently been found to correlate with the routes taken by individual birds from Europe to their African wintering grounds. However, the genomic location of this region remains unknown. Here, we characterized MARB using a combination of approaches to understand how it evolved. We describe the region using long-read genome assemblies of two willow warbler subspecies (*P. t. trochilus* and *P. t. acredula*), two related species, the common chiffchaff (*P. collybita*) and the greenish warbler (*P. trochiloides*), and whole genome sequencing data from 76 willow warblers. Finally, we applied karyotyping and fluorescent in situ hybridization techniques on willow warbler spermatocytes to cytogenetically locate MARB. Due to the many repeats, we cannot order scaffolds *in silico*, but probe hybridization on the karyotype shows that MARB constitutes a single locus (~27.5 Mb) spanning most of the 11^th^ largest chromosome in the willow warbler genome. Interestingly, the MARB regions of all species share several characteristics such as relatively high GC content (50%), a high density of specific repeat families and notably, more than 800 olfactory receptor sequences. Regions homologous to MARB may exist in several migrant bird genomes, though currently unassembled due to their complexity. Resolving these in species with similar migratory polymorphisms to willow warblers will be essential to determine whether MARB influences migratory behaviour across species.

## Background

Bird migration is a fascinating phenomenon, which has evolved to fulfil different needs across the annual cycle and it has been noticed by mankind for thousands of years, with numerous references to the periodic movement of birds since the time of Aristotle (1). Yet, it is still unclear how some birds can leave their breeding grounds at a specific time and tirelessly fly in a particular direction across mountains, large water bodies or deserts, in solo flights. How is a bird sure about the correct orientation? What pushes the bird’s decision of when to start and end its journey? About half of the bird species in the world are described as migratory (2) and display a wide variety of routes and distances (3). This complex behaviour is a syndrome, characterized by a suite of integrated traits encompassing behavioural, physiological and genetic factors (4).

It is well established that birds rely on a genetic endogenous program to complete the migratory cycle (5–7) to a greater or lesser degree. Such genetic determination is predominant in solitary migrants, usually passerines, which typically fly at night (8,9). Due to the exponential improvement of sequencing technologies and miniature tracking devices, numerous contributions have been made to the field of genetics of bird migration in the past years (10–13). Several studies have found correlations between genes or genomic regions and different migratory traits such as distance (14), timing (15) and wintering locations (16). The genetic basis seems to be quantitative (17) and recent research has proposed a diversity of regulatory mechanisms behind some of these traits (18). However, our understanding of the genetic underpinnings is still very poor.

One of the most enigmatic traits shaping migratory routes is orientation, as juvenile birds can find population-specific wintering grounds without any guidance from adults. Until now, few studies have shown associations between migration direction and any genomic region. The strongest correlation to date has been described in the willow warbler (*Phylloscopus trochilus*). Two subspecies (*P. t. trochilus* and *P. t. acredula*) occur in Europe and meet at migratory divides in central Scandinavia, East of the Baltic Sea (19) and in the Åland archipelago (20). Whereas *P. t. trochilus* migrates through Western Europe to West Africa, *P. t. acredula* takes an eastern route across the Balkans to East and Southern Africa (21). Comparative genomic analyses and geolocator tracks from pure subspecies and hybrids showed that a repeat-rich region named migration-associated repeat block (MARB) explains 64% of the natural variation in the migratory direction of two migratory phenotypes (13).

MARB was initially described in Caballero-Lopez et al. (22) as a region comprising seven scaffolds (with a total length of 22.8 Mb) in the *P. t. acredula* genome, which carries a diagnostic transposable element (TE) from the LTR family named WW2-derived. Notably, qPCR analyses showed that eastern migrating birds (subsp. *acredula*) carry more copies of this TE (7–45) than birds following the western flyway (0-6; subsp. *trochilus*). The copy number variation of WW2-derived was sampled in two willow warbler families in the hybrid zone between *P. t. trochilus* and *P. t. acredula* in central Sweden. This showed that the copies appeared to be inherited as intact blocks, consistent with a Mendelian inheritance pattern (Table S5; 13). The MARB region has a higher GC content (50%) than the rest of the genome (43%) and consists of various repeats, including olfactory receptor (OR) genes and a high density of TEs (22). Despite combining long-read sequencing methods, linking MARB to any single-copy genomic region with known chromosomal locations has not been possible (23).

The presumably strong effect of MARB on migration direction calls for a more detailed characterization of this region (Figure 1). To understand how MARB has evolved several questions arise: 1) Do the scaffolds identified in *P. t. acredula* (MARB-a) have an orthologous region in the *P. t. trochilus* genome (MARB-t)? 2) Is the MARB region unique to willow warblers or does it exist in other *Phylloscopus* species? 3)What is the location of MARB in the genome and is it linked to a region with single-copy genes and/or a known chromosomal location?

**Figure 1.**
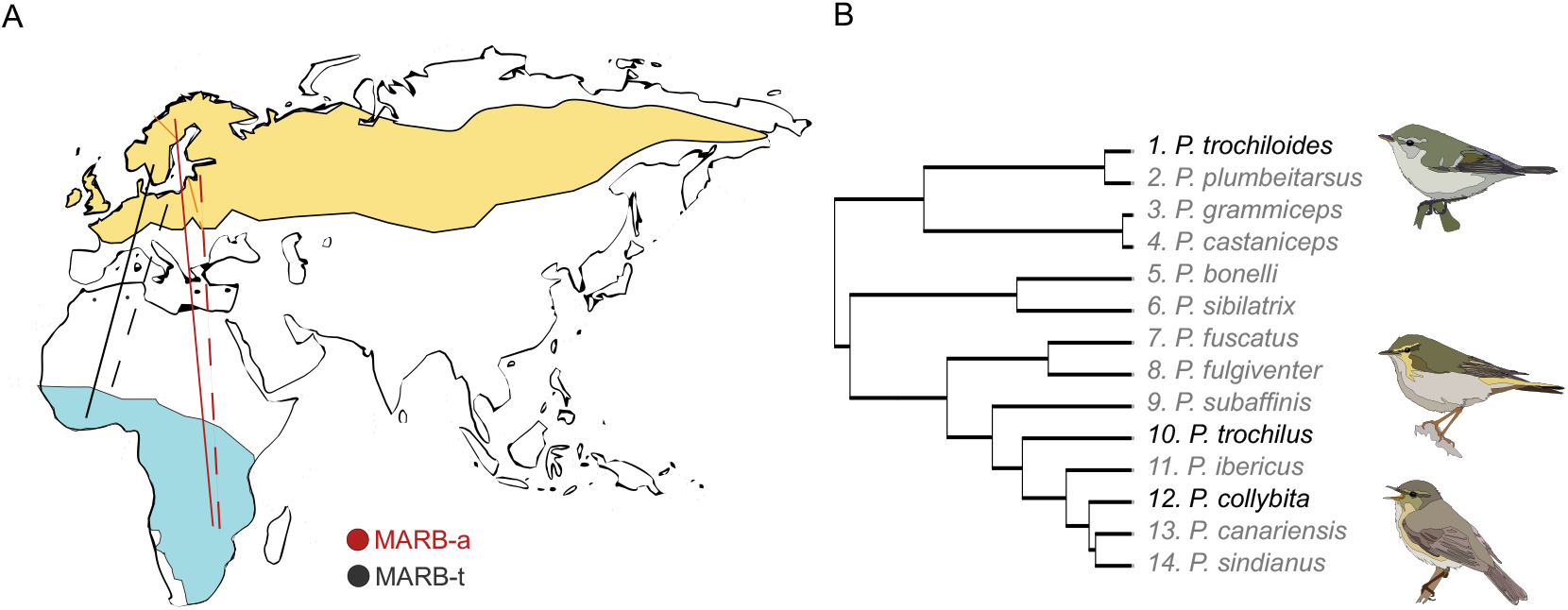
Study of the MARB region across three *Phylloscopus* species. **A)** Migratory directions of willow warblers between breeding (yellow) and wintering (blue) grounds. Routes depicted for birds containing MARB-a (red lines) and MARB-t (black) across both migratory divides (orange lines). Continuous lines summarise published data (13); dashed lines summarise unpublished data. **B)** Species from the genus *Phylloscopus* in which we describe MARB. The split of greenish warbler from the willow warbler and chiffchaff clade is estimated to have happened c.a., 11 million years ago (32).

To address these questions, we analysed high-quality genome assemblies of both willow warbler subspecies and two related species in the genus *Phylloscopus:* the common chiffchaff (*P. collybita* ssp. *collybita)* and a more distant relative, the greenish warbler *(P. trochiloides* ssp. *viridanus)*. In parallel to the genomic analyses, we utilized fluorescence *in situ* hybridisation (FISH) to visualise MARB on willow warbler (*P. t. trochilus*) meiotic karyotypes.

## Results

### MARB in willow warblers

The resequenced individuals were grouped into birds with MARB-a (n = 38) and without (n = 38) based on the established qPCR assay (22). The reads mapped to the *trochilus* reference genome, identified 11 scaffolds (total length 44.0 Mbp) with strongly elevated F_ST_ values between these two groups (Figure S1). These scaffolds have an elevated % GC compared to the rest of the genome (50.7 versus 43.8), a 40-fold higher density of olfactory receptors, and specific repeat families. These sequence characteristics are shared with the seven scaffolds in the acredula genome previously identified as MARB based on the presence of the WW2-derived TE (Table S1).

An MDS plot for the SNPs within these eleven scaffolds generated three clusters, presumably corresponding to three genotypes and supporting a single locus (Figure S2). In support, the qPCR estimated copy number of the WW2-derived TE was strongly correlated (r = 0.834, P = 10^−22^) with the first component of MDS.

The total length of the MARB-t scaffolds (approx. 44.0 Mbp) was longer than the total of the previously identified MARB-a scaffolds based on the presence of the WW2-derived TE (approx. 23.0 Mbp). We therefore investigated the possibility that there are MARB-a scaffolds without WW2-derived TEs. To verify this, we mapped the reads of the 76 individuals to the *acredula* genome and identified seven more scaffolds with strongly elevated F_ST_ values between these two groups. These scaffolds also share a similarity of repeat content to the other scaffolds classified as MARB, bringing the total length for MARB-a to 35.7 Mbp (Table S1).

Though the MARB-t and MARB-a scaffolds share many sequence characteristics and likely represent alternative haplotype groups of the same locus, they align poorly and only over fragmented stretches (Figure S3). The strongly repeated structure has probably promoted non-allelic homologous recombination (NAHR; 24) within MARB-t and MARB-a, respectively, thus shuffling around various repeat blocks, leading to the divergent organisation of MARB in the two subspecies. Moreover, we cannot exclude the possibility of highly divergent haplotypes of MARB within *trochilus* and *acredula*, respectively, which could have been erroneously categorised as independent scaffolds, instead of alternative haplotypes, in the assemblies. Hence, the total length of MARB might be shorter than the sum of the lengths of all scaffolds.

The repetitive structure of MARB may suggest that it has expanded by successive duplications. If that is the case, we predict that the olfactory receptors of MARB are more closely related to each other than those in the rest of the genome. We used the more complete *trochilus* genome to address this question. After filtering, we identified 1402 OR sequences. These belonged to the clades α (n = 3), γ (n = 8) and γ-c (n = 1391), the latter strongly dominating as in other bird genomes (25). In contrast to our expectations, we found that the MARB ORs were not restricted to specific clusters in the phylogeny but were highly mixed with ORs from the rest of the genome (Figure 2). Notably, 57% of the predicted OR genes in the *trochilus* genome are located in MARB-t. Furthermore, one end of Scaffold97 is flanked with telomeric repeats (TTAGGG × 237), suggesting this spans the chromosome end. In the case of the *acredula* genome only 20% of ORs are located in MARB-a scaffolds, but this is likely due to the large regions of unknown nucleotides (N’s) constituting 39% of the MARB-a scaffolds (Table S1).

**Figure 2.**
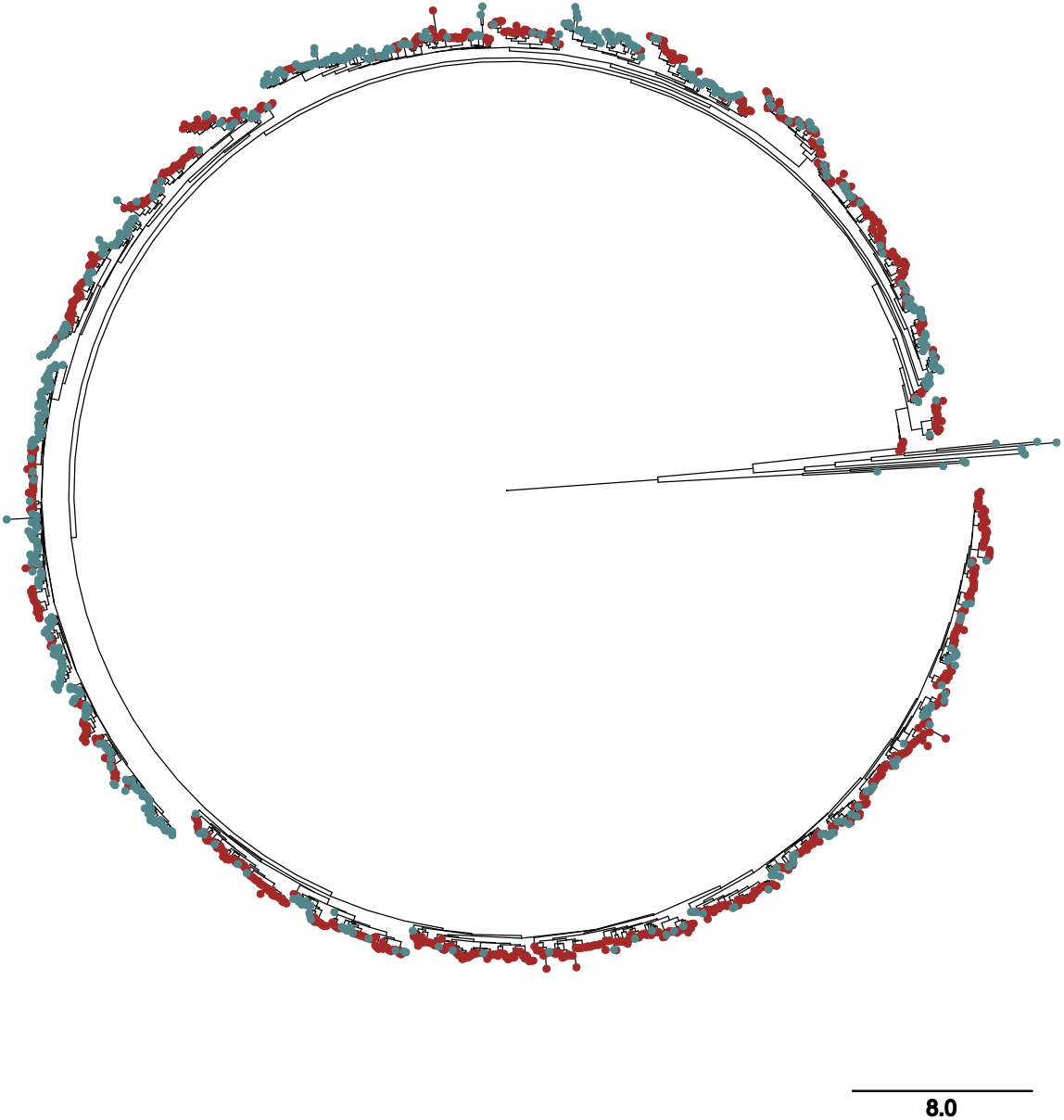
Phylogeny of 1,402 olfactory receptor (OR) genes from the willow warbler *trochilus* assembly, located in MARB (red) and non-MARB (blue) scaffolds. The majority of genes (n = 1,391) belong to the γ-c clade and are distributed throughout the tree without apparent sub-structuring relative to genomic location. All genes in clade α (n = 3) and γ (n = 8) were located on non-MARB scaffolds. Ten chicken non-OR genes were used as an outgroup and subsequently collapsed for visualisation purposes. The scale bar represents the expected number of substitutions per site.

### MARB in chiffchaffs and greenish warbler

We used BLAST of MARB segments (approx. 5 kb) from *trochilus* and *acredula* to identify candidate MARB scaffolds in the genomes of *collybita* (MARB-c) and *viridanus* (MARB-v). Among scaffolds with repeated hits, we identified six *collybita* (total length 28.0 Mbp) and seven *viridanus* scaffolds (total length = 46.8 Mb) that also shared several other willow warbler MARB characteristics: a GC content of ~50% and elevated abundance of ORs and certain repeats (Table S1). Most ORs in the *collybita* and *viridanus* genomes (72%) are located in the identified MARB scaffolds. Four of the scaffolds in *collybita* contained the WW2-derived TEs, arranged in a reversed direction with a copy of WW2 ancestral TE as previously described for *acredula* (22) further supporting their classification as MARB. With few exceptions (Figure S4), most parts of the MARB in *trochilus* and *collybita* failed to produce any longer alignments. Telomeric repeats flank two *collybita* (#32 and #54) and one *viridanus* (#39) scaffolds.

### Meiotic karyotypes

Spermatocytes showed a consistent number of 40 bivalent chromosomes through immunostaining (Figure 3). This results in a total of 80 chromosomes for willow warblers plus one extra univalent germline restricted chromosome (GRC) that is labelled across its length by CREST antibodies (Figure S5). The chromosome pairs could be ordered by size, with the 10 largest chromosomes being approximately 6.26 µm long and the rest progressively shorter. However, the boundary between macro and microchromosomes is not clearly defined.

**Figure 3.**
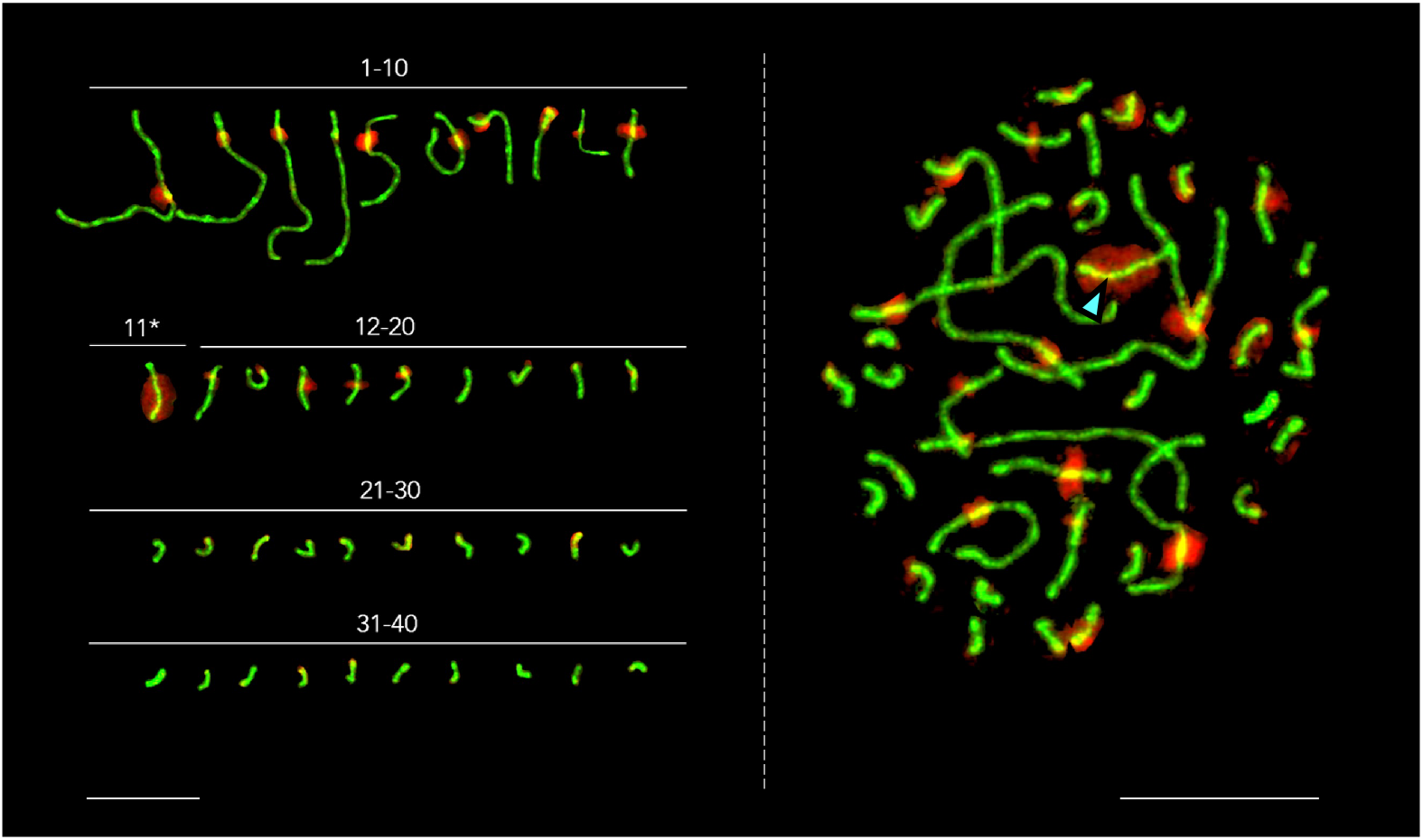
*In situ* hybridization results showing the location of MARB. On the right, spermatocyte (spread from willow warbler (*P. t. trochilus*) testes immunostained with antibodies against synaptonemal complex lateral elements, SYCP3, (green), and a PCR-synthesized probe for MARB (red) hybridising almost entirely to one bivalent (blue arrow) and to (peri)centromeric regions of most bivalents. On the left, the chromosomes are sorted by size, showing that the MARB chromosome is the eleventh longest in the genome (labelled as 11*). Scale bars = 10 µm.

The estimated size of the MARB region across the four genome assemblies varies (*acredula*: 36 Mbp; *trochilus*: 44 Mbp; *collybita*: 28 Mbp; *viridanus*: 46.8 Mbp). We cannot discard that such measurements are either an overestimation resulting from the concatenation of redundant scaffolds, or an underestimation due to some regions not being classified as MARB. The total length of the full willow warbler genome is ~1300 Mbp, corresponding to ~215.6 µm (SD = 15.6 µm; n = 10) in the karyotype. The most extensive probe binding occurs across 4/5 of the length of the 11^th^ longest chromosome (average length = 5.71 µm; 34.42 Mbp) identifying it as the chromosome containing the MARB region (Figure 3).

Our probe also hybridises with elements in the proximity of centromeres (Figure 3), confirmed by mitotic metaphase stage spreads (Figure S6). We have not been able to confirm that the full-length probe sequence (~1800 bp) matches any centromeric regions in the *trochilus* genome assembly. However, this does not conflict with the result of the hybridization experiment for the following reasons: first, nearly all chromosomes in the genome assembly have gaps over the presumed centromeric regions. Hence, we do not know the complete sequence of most centromeres or perhaps even pericentromeric regions. In addition, these regions typically have a high and variable repeat content (26) that could have caused unspecific binding to the probe, given that it had to be fragmented before hybridisation.

## Discussion

Here we confirm that a repetitive region, which displays the highest association with migratory direction to date (13) and was initially identified in *P. t. acredula* (22) as MARB-a is present as an homologous region in *P. t. trochilus* as MARB-t. Remarkably MARB is so divergent across taxa that it cannot be aligned but similarities in GC content (around 50%), the abundance of olfactory receptor sequences and specific repeat families also confirm homology in *P. c. collybita* as MARB-c and *P. t. viridanus* as MARB-v. Using karyotyping and FISH techniques on willow warblers (ssp. *trochilus*) we unravelled the chromosome location of MARB to the 11^th^ largest chromosome (34.4 Mbp) with an estimated size of 27.5 Mbp. A multidimensional scaling analysis of SNPs in 76 resequenced willow warblers (encompassing pure subspecies and hybrids) reveals three distinct MARB clusters presumably corresponding to the three genotypes (two parental homozygotes for MARB-t and MARB-a and one heterozygote), further supporting that MARB is a single locus.

It is worth noting that the karyotype-based size estimate of MARB falls somewhat on the short end of the length range estimated from our genomic analyses of this region across the four *Phylloscopus* taxa (26-46.8 Mbp) and the total length of four MARB-like scaffolds (47.2 Mbp) in a recent genome assembly in *P. t. trochiloides* (27). This disagreement between cytogenetic and bioinformatic estimates can be due to a few reasons: though the MARB regions in these taxa show similar types of repeats, they fail to align over longer stretches, even between the willow warbler subspecies (MARB-a and MARB-t). This can be explained by the high repeat content facilitating frequent unequal recombination events, thereby promoting diversity and reducing synteny (28). The largest MARB might be an overestimation resulting from the concatenation of redundant scaffolds due to the high heterozygosity (29). Conversely, other scaffolds containing segmental duplications might have been misinterpreted as one locus (30). Lastly, high TE content is a known driver of genome size variation across individuals (31). Therefore, we cannot discard individual variation in the size of MARB, as the genome assembly and the karyotype originate from different birds.

Within the genus *Phylloscopus*, the greenish warbler diverged from the willow warbler and the chiffchaff about 11 mya (32). The presence of MARB in these species indicates that this large region existed already in the common ancestor to all *Phylloscopus*. While we cannot entirely rule out the possible assembly errors mentioned above, the observed size variation among these species suggests that chromosomal contractions or expansions could be more common than we thought. In line with this, while birds generally maintain a highly conserved chromosome number (2n=78-82; 33) notable differences in chromosome size and morphology, even between closely related species, indicate that such structural variation may be more frequent than the stable chromosome count implies (34).

The MARB genotype is predicting the migratory direction of willow warblers with remarkable precision (13). However, the mechanism by which MARB influences this trait remains an enigma. Regulatory differences affecting the underlaying genes are thought to be important drivers of variation in migratory phenotypes between (35,36,18) and within (15) populations. Concerning MARB, this is a plausible scenario given its characteristics; high repeat density, GC content and lack of single-copy genes may indicate that this region is heterochromatic and silenced (37), though it may still interact with other genomic regions. Heterochromatin is involved in the spatial organization of chromosome regions in the nucleus and can potentially affect gene expression elsewhere (38). In the case of willow warblers, the epistatic interaction between MARB and chromosome 1 on the migratory behaviour (13) could point to a process through which MARB-a/MARB-t act as regulatory switches for migratory direction.

Because of the repeated structure, the MARB-a and MARB-t haplotypes align poorly which makes it challenging to identify regions exclusively differing between them. However there is one consistent difference between eastern and western migrants: the copy number variation (CNV) of the 5-bp indel within the LTR-transposon sequence which aided the original identification of the MARB haplotypes, known as WW2-derived (22). This poses the question of whether this TE copy number has functional relevance in determining migratory direction. There is evidence for differential gene expression linked to retrotransposon CNV in other organisms where they are involved in oncogenesis, senescence and transcriptional levels (39,40). Given the absence of coding genes in the WW2 surroundings, their hypothetical effects could be to act as enhancers of more distant genes (41) or differences in chromatin folding among other possibilities (42,43).

Another noteworthy characteristic of MARB is its gene composition. At least 50% of the OR genes total (>1000, mostly from clade γ-c) across the three *Phylloscopus* species are located in the MARB scaffolds (3% of the genome). This amount may seem excessive compared to most bird genome assemblies that contain <100 OR genes (25,44,45). However, in high quality PacBio genome assemblies of two additional species, both migratory and distantly related to the *Phylloscopus* warblers, >1000 ORs have been detected: the northern flicker *Colaptes auratus* (GCA_015227895) and the purple martin *Progne subis* (GCA_022316685) (45). In the flicker, the ORs are clustered in a large unplaced scaffold (JAAWVA010000030; 36 Mbp) whereas purple martin has the ORs scattered in >100 smaller unmapped contigs (< 1 Mbp).

This large variation in OR gene numbers across birds may suggest a rapid expansion of ORs in some lineages. Alternatively, and supported by current research on avian olfaction, the number of OR genes may be underestimated in many species. If these genes mainly occur in repeat-rich regions they will be missing from genome assemblies based on short reads. For example, the previous genome assembly (GCA_016584745.1_LUNI_Phtr_3; 10x chromium Illumina HiSeqX) of the same *trochilus* sample as used in the present analyses contained only 386 ORs (45). Likewise, the number of ORs in the chicken *Gallus gallus* genome assemblies has increased with the improved techniques of sequencing (272 in Ggal4, 355 in Ggal6, 426 in GGswu1; International Chicken Genome Consortium, (46). Interestingly, 386 of the OR’s in GGswu1 are located on chr29, one of the so-called dot chromosomes. Likewise, in the most recent assembly of the zebra finch (*Taeniopygia guttata*; bTaeGut7), a micro-chromosome (chr30) contains most of the OR’s in the genome (47).

Apart from the OR content, there are other hints that MARB might be homologous to the chicken dot chromosome chr29. This chromosome has a length approximately 6.4 Mbp and the region starting at 2.8 Mbp shares key characteristics with MARB, such as a high density of certain repeat families (i.e., Rnd-3_family-52, Rnd-3 family_21) and an equally high GC content (~50%). Interestingly, the region up to 2.8 Mbp is described as euchromatic with several single copy genes and an exceptionally high GC content (>60%) which potentially hindered the sequencing of the *Phylloscopus* with the PacBio technique (46). However, the chicken chr29 has the highest map score to chr31 in the zebra finch genome, and not to the most OR rich chromosome (chr30).

Hence, the genome assemblies of the chicken and the zebra finch do guide us to the presumed single-copy region linked to MARB, represented in our karyotype by the portion of the 11^th^ largest chromosome where the probe did not bind. This region could contain the gene(s) which together with MARB determine migratory direction in willow warblers (13).

Olfactory information is processed in the piriform cortex, in tight connection with the hippocampus, an area directly involved in short and long-term memory and learning (48). If there is indeed ongoing gene expression within MARB, it is exciting to hypothesize a role of olfaction in migratory behaviour, given that reliance on olfactory maps has been observed across several bird species in the wild (49–51). A strong sense of smell may help these birds memorize their route to wintering grounds and back. It is difficult, though, to imagine which role ORs would play in a juvenile bird that has never migrated before, beyond gathering new information. It seems more likely then, that the high OR abundance in MARB supports navigation, rather than determining migratory directions.

## Conclusions

Less than 10% of passerine species have been karyotyped (33,34). Even migratory species that are represented by high-quality genome assemblies such as blackcaps (*Sylvia atricapilla*) and pied flycatchers (*Ficedula hypoleuca*) lack karyotypes. Such genome assemblies are still unable to resolve high-complexity regions, which, although often treated as junk DNA (52), can underpin complex behaviours such as the migratory syndrome. We show that in several (and potentially all) *Phylloscopus* species MARB contains a much higher amount of intact OR copies than the average number found in most bird genome assemblies. Although transcription remains to be demonstrated, the integrity of the sequences hints towards functional relevance. This suggests that the sense of smell in birds may have been repeatedly underestimated (53) and could influence migratory phenotypes.

Notably, OR-rich chromosomes (or large scaffolds) are absent from assemblies used in genotype-phenotype association studies of bird migration (45). For example, the genomes of the common blackcap *Sylvia atricapilla* (GCA_009819655.1), the Swainson’s thrush *Catharus ustulatus* (GCF_009819885.2) and the myrtle warbler *Setophaga coronata* (GCA_001746935.2) used in the study of migration in *Vermivora* warblers (16) lack such regions. This raises the possibility that in such genomes, OR-rich chromosomes were not assembled, rather than simply absent. We therefore expect that higher-quality bird genomes will reveal that more species have particularly OR-rich chromosomes. It will be essential to evaluate whether these are homologous to MARB and, further, to test whether they are associated with migration differences in other species.

This study provides a valuable contribution to the understanding of the structure and evolution of genomes. It highlights the importance of combining karyotypes and high quality genome assemblies to resolve and characterize challenging regions that influence complex phenotypes. We raise the question of whether the effect of MARB on migratory behaviour crosses the species boundaries, not only in other *Phylloscopus* taxa where it is present, but also beyond.

## Methods

### Reference genomes

We used the long-read genome assemblies of four individuals representing four taxa of *Phylloscopus* warblers. Three genomes, *P. t. trochilus* (male), *P. t. acredula* (male), and *P. c. collybita* (female), are described in a previous publication (23) and are available at NCBI under bioproject PRJNA550489. The assembly of *P. trochiloides viridanus* (male) was generated for the present study. In the following paragraphs, we refer to these taxa as *trochilus, acredula, collybita* and *viridanus*, respectively.

Briefly, the genome assembly of *acredula* was generated from a combination of continuous long reads (CLR, Pacific Biosciences, CA, USA) generated on eight SMRT (Single Molecule Real-Time) cells on a Sequel platform (PacBio), linked reads (10X Genomics, CA, USA) and optical mapping data (BioNano Genomics, USA). The genomes of *trochilus* and *collybita* were assembled using High-Fidelity reads (HiFi, PacBio) originating from two cells of Sequel II (PacBio) for each species and the *trochilu*s assembly was additionally scaffolded with BioNano data. DNA for the *viridanus* assembly was collected from an adult male captured at Juodkrante (55.54°N, 21.12°E), Lithuania, during the 2022 breeding season. Approximately 30μl of blood was drawn from the brachial vein and stored in pure ethanol at ambient temperatures for one week, and after at −20°C until extraction of DNA using a Nanobind extraction kit (Circulomics, MD, USA). The sample was sequenced on two cells on a Revio (PacBio) using a HiFi setup, which generated 6,812,139 reads with a mean length of 13,240 bp. As with the previous three genomes, extraction of DNA and sequencing was performed at SciLifeLab, Uppsala, Sweden. The HiFi reads were assembled into 555 contigs using Hifiasm version 0.19.5 (54) with default settings.

The improved techniques for long read sequencing from *acredula* to *viridanus* explain why the genome assemblies contain a progressively lower number of unresolved regions (Table S1).

The willow warbler genomes were annotated for protein-coding genes using a gene predictor together with RNA-seq data and protein data from other species as described in Lundberg et al. (23). To facilitate genome comparisons, we used the annotations from *trochilus* to annotate the *collybita* genome assembly. The *viridanus* genome was annotated using Augustus (55) with the “chicken” training set. The repeats were annotated in all genomes with RepeatMasker (56) using a library that contains both *de-novo* and publicly available bird-specific repeats (23).

### Resequencing data

All libraries for genome resequencing were prepared from 1μg DNA using the TruSeq PCRfree DNA sample preparation kit (cat#20015962/3, Illumina Inc.) targeting an insert size of 350bp. The library was prepared according to the manufacturer’s instructions (guide#1000000039279). For details see Table S2.

### MARB sequences identification

We used 76 resequenced willow warbler males, mainly from the migratory divides (in Sweden, Poland and Lithuania) to examine sequences potentially representing MARB in *trochilus* (MARB-t) and *acredula* (MARB-a). If MARB is a single locus region, as previously suggested (13); Table S5), we expect that MDS analyses of SNPs located on MARB-scaffolds should result in three clusters corresponding to MARB-t homozygotes, heterozygotes (MARB-t / MARB-a), and MARB-a homozygotes. Furthermore, these three genotypes are expected to show different WW2-derived TE copy numbers as scored by the qPCR assay (22), with MARB-t homozygotes having <7 copies, heterozygotes having >7 but, on average, fewer copies than MARB-a homozygotes.

We identified MARB-like scaffolds in the genomes of *collybita* and *viridanus* by searching for scaffolds with a content similar to willow warbler MARB. The key characteristics were 1) the presence of the WW2-derived TEs, 2) a high abundance of ORs and other MARB-enriched repeats, and 3) a GC content of approx. 50% (Table S1). We also inspected the flanking region of these scaffolds for the presence of single-copy genes that could potentially link them to annotated chromosomes.

### Mapping and variant calling

The reads were mapped to the *trochilus* and *acredula* assemblies using bwa mem (v0.7.17; 57) with default parameters. The mapping was done independently for each lane. The resulting alignments were sorted using SAMtools (v1.9;58). Duplicates were then identified and marked using the Picard tool (https://broadinstitute.github.io/picard/) MarkDuplicates (GATK v4.1.4.1), and alignments from all lanes were combined.

We conducted variant calling using freebayes (v1.3.2; flags: --standard-filters --min-alternate-count 5 --min-alternate-fraction 0.2 --use-best-n-alleles 3 --skip-coverage 3200; 60). The raw variants were first filtered using vcffilter in vcflib (v1.0.1) with the following criteria: quality score > 30, at least one alternate observation on each strand (SAF > 0 & SAR > 0) as well as at least one read balanced to the right and to the left (RPR > 0 & RPL > 0). We performed subsequent filtering steps with vcftools (v0.1.14; 61). Variants with low (<10) or high (>68) mean sequencing depth were removed, and genotypes with low depth (<4) were set as missing. Next, variants with high missingness (>0.5) and those overlapping annotated repeats were removed. Finally, only bi-allelic variants were retained for analyses.

### Multidimensional scaling analysis

Willow warbler VCF files were converted to PLINK format using PLINK2 (v2.00-alpha-2.3-20200124; 62). Variants with minor allele frequency (MAF) < 0.1 were excluded and linkage-disequilibrium (LD) pruning was performed with PLINK (v1.90b4.9) and parameters --indep 50 5 2. PLINK was then used to run multidimensional scaling (MDS) calculations on the LD-pruned set.

F_ST_ was calculated between two sets of birds using VCFtools (v0.1.14), both for individual SNPs and in windows of 10 kb. For the windowed calculations (only variants with MAF > 0.1 were included), and only windows containing at least 25 SNPs were kept.

### Analyses of olfactory receptor genes (OR)

We predicted “intact OR genes” if they passed the following filtering steps: 1) absence of a stop-codon in the sequence, 2) a length of the protein sequence > 250 amino acids (44), 3) seven transmembrane domains predicted by DeepTMHMM (62), (4) a higher similarity to known chicken OR genes than to (non-OR) G-protein coupled receptor (GPCR) (25). For this filtering step, we used the non-OR chicken protein sequences from Lagerström et al. (63) and built a phylogenetic tree with the predicted OR genes plus the chicken non-OR CPGR genes with IQ-TREE (version 2.2.2.2; 58) with the parameter “-m LG”. Any predicted OR genes that were phylogenetically more closely related to the non-OR GPCRs were removed.

The protein sequences that passed the filtering were aligned in Clustal Omega (version 1.2.4; 59) and a phylogenetic tree was built using IQ-TREE (version 2.2.2.2), with the options -m MFP -B 1000 -alrt 1000, assessing branch supports with both SH-aLRT and the ultrafast bootstrap approximation (66) and ModelFinder (67) used to identify the best-fit substitution model.

### Sample collection for karyotyping

Three adult male willow warblers were captured with mist-netting techniques and playback in their established territories, blood-sampled and euthanised shortly afterwards, during the 2024 breeding season. The trapping was performed in Czechia (Dolní Počernice; 50°05’51.6”N 14°34’47.6” E) within the *P. t. trochilus* distribution range. Testes and bone marrow were dissected right away after euthanasia by cervical dislocation at the Faculty of Science, Charles University (Prague).

### Mitotic chromosome preparation

Bone marrow was flushed out of the tibia using a syringe needle with Dulbecco’s Modified Eagle’s Medium (Sigma Aldrich), enriched with 10% (v/v) foetal bovine serum (Biowest), 3% (v/v) phytohaemagglutinin (GIBCO), 1% (v/v) L-glutamine (Sigma-Aldrich), 1% (v/v) penicillin/streptomycin solution (GIBCO) and 1% (v/v) lipopolysaccharide (Sigma Aldrich). After a 3h incubation at 37 °C in 6 ml of medium with 75 µl of 0.1% colcemid solution (Roche), the cells were transferred to pre-warmed hypotonic buffer (0.075M KCl) for 30 minutes at 37°C. Subsequently, they were washed 3 times with 5 ml of cold fixative solution (methanol: acetic acid, dilution 3:1 ratio).

### Meiotic chromosome preparation and immunostaining

We prepared synaptonemal complex (SC) spreads from freshly dissected testes from the three individuals, and followed Poignet et al. (68) with slight modifications. Briefly, testes were cut into two pieces to release germ cells, and kept in hypotonic buffer (30 mM Tris, 50mM sucrose, 17 mM trisodium citrate dehydrate, and 5 mM EDTA; pH 8.2) for 40 minutes. Then, they were mashed into a 100 mM sucrose solution. Solids were removed with tweezers and the cell suspension was applied to a pre-treated slide with 1%PFA and 0.15% Triton X-100 (Sigma Aldrich). The slides were then placed in a humid chamber for 90 minutes, washed in 1× PBS for 2 minutes and directly used for immunostaining.

We conducted immunostaining before probe hybridisation to observe the SC in spermatogenic cells. Briefly, we incubated the slides in a wet chamber for 3 hours at room temperature with rabbit anti-SYCP3 polyclonal primary antibody (dilution 1:100, Abcam, ab15093), which binds to the lateral elements of the synaptonemal complex. Then, the slides were washed three times at room temperature in 1× PBS and incubation was repeated with a secondary antibody (anti-rabbit IgG-AlexaFluor488, dilution 1:200, A-11008, Invitrogen). All antibodies were diluted in PBT (3% BSA and 0.05% Tween 20 in 1× PBS) before use. Afterwards, the slides were rewashed and dehydrated in an ethanol series (70–80–96%, 3 minutes each). When they dried, they were used for probe hybridization straight away.

### MARB probe synthesis and fluorescence in situ hybridization (FISH)

The MARB-binding probe was designed in Geneious Prime (v 2024.0.3) using the multiple alignment of a repeat combination (Rnd-5_family-1399 + Rnd-3_family-21 + Rnd-4_family-564 + Rnd-3_family-21) that is particularly abundant in willow warbler MARB scaffolds compared to the rest of the genome. The sequences used in the alignment were found by blasting the repeat combination to the reference genomes of the two willow warbler subspecies and selecting 20 divergent hits from each. It was synthesized and labelled with digoxigenin-dUTP (Roche) by PCR amplification (50 µl reaction) from genomic DNA extracted from the blood of a *trochilus* bird following the ammonium acetate protocol (69). The primers used were MARB_1aF (5’-GTCAGGGACCTGCAGCTTG-3’) and MARB_1aR (5’-CCATCTGCTCTGCTGAGGATG-3’): 95°C for 5 minutes as initial incubation, and 35 thermal cycles (95°C for 1 minute, 59°C for 30 seconds, 72°C for 1 minute) followed by 72°C for 6 minutes end phase. The PCR product was then purified using the Gel/PCR DNA Fragments Extraction kit (Geneaid) according to the manufacturer’s protocol. The purified PCR product was approx. 1800 bp, long and 1 µg was subsequently treated with nick-translation (Nick-translation kit, Abbott) in a 50 µl reaction for 104 minutes at 15°C to shorten the probe to a final size of approx. 200-300 bp. The probe was then precipitated over two days at −20°C with 5 µl salmon sperm DNA (10 mg/ml, Sigma Aldrich), 5 µl of 3M sodium acetate (pH 5.2) and 200 µl of ice-cold 96% ethanol. After precipitation, the probe was washed with ice-cold 70% ethanol, and the dried pellet was resuspended in 60 µl of hybridization buffer (50% formamide, 2× SSC, 10% dextran sulfate, 10% sodium dodecyl sulfate and 1× Denhardt’s buffer, Ph 7.0).

The probe was denatured at 86°C for 6 minutes and then kept at −20°C until use (~10 minutes). Meanwhile, the pachytene spreads were denatured in 75% formamide/2 × SSC at 75°C for 3 minutes and dehydrated in an ice-cold ethanol series (70-80-96%, 3 minutes each). After drying, 20 µl of the probe (325 ng) in hybridisation buffer was placed on each slide and hybridised for 48 h at room temperature in a humid chamber. Post-hybridisation washes were performed on the slides in 0.2 × SSC at 50°C three times before incubation with anti-digoxigenin-rhodamine (red) at room temperature for 3h in a humid chamber. The slides were then subsequently washed with 4 × SSC/0.01% Tween 20 (Sigma Aldrich) at 44°C, dehydrated through ethanol series (70-80-96%) and dried. Finally, 20 µl of Vectashield/DAPI (1.5 mg/ml; Vector Laboratories) was added for counterstaining of the chromosomes.

### Microscopy and image processing

Meiotic spread images were acquired and examined using an Olympus BX53 fluorescence microscope with a DP30BW digital camera (Olympus). We inspected approx. 300 pachytene-stage spermatocytes, from which the 10 best spread nuclei were selected for measurements analyses and repeatability. SC length was measured in triplicate to reduce technical error using ImageJ2 (v 2.14.0/1.54f). Processing of the images, including background removal and arrangement of SC by size was done using Adobe Photoshop CS software (Adobe Systems).

## Declarations

### Ethical permits

Sample collection was performed following regulations in accordance with institutional guidelines. The work with birds in the Czech Republic was approved by Prague City Hall, Department of Nature and Landscape Conservation (permission no. MHMP 950972/2024).

## Availability of data and materials

All resequence and genome data will be made available at ENA (European Nucleotide Archive; https://www.ebi.ac.uk/ena/browser/home) upon publication.

## Funding

The cytogenetic work was supported by the Czech Science Foundation grant to RR and DD (23-07287S). Sequencing was performed by the SNP&SEQ Technology Platform in Uppsala. The facility is part of the National Genomics Infrastructure (NGI) Sweden and Science for Life Laboratory. The bioinformatic analysis was supported by the SciLifeLab & Wallenberg Data Driven Life Science Program, Knut and Alice Wallenberg Foundation (grants: KAW 2020.0239 and KAW 2017.0003), and by the National Bioinformatics Infrastructure Sweden (NBIS) at SciLifeLab. The computations were enabled by resources provided by the National Academic Infrastructure for Supercomputing in Sweden (NAISS), partially funded by the Swedish Research Council through grant agreement no. 2022-06725. The rest of the study was supported by grants to VCL from the Royal Physiographic Society of Lund, and the Jörgen-Lindströms foundation, and to SB from the Swedish Research Council (2021-03853), Erik Philip-Sörensens stiftelse (G2019-029, G2022-014), National Geographic Society (WW-208R-17) and Olle Engkvists Stiftelse (232-0112).

## Author’s contributions

Conceptualization: V.C-L., S.B. Field data collection: O.K., J.R., V.C-L., S.B., Laboratory work: V.C-L., D.D. Laboratory work support: L.O-H, R.R, V.T., M.A. Bioinformatic analyses: D.E., E.P-W., M.Lu. Writing: V.C-L. Review and editing: all authors.

## Acknowledgements

We are grateful to Carlos Lara, Eleonora Pustovalova and Tomáš Pavlica for the valuable advice throughout the karyotyping labwork.

## Notes

### Competing Interest Statement

The authors have declared no competing interest.

