## Supplemental material for "A songbird karyotype: cytogenetic confirmation of a migration-associated region rich in olfactory receptor genes"

### Supplementary material for: “A songbird karyotype: cytogenetic confirmation of a migration-associated region rich in olfactory receptor genes”

#### Methods

*Mitotic chromosome preparation*

The cells were spread on slides using the dropping technique and stained with Giemsa solution (5% Giemsa in 0.07 M phosphate buffer, pH 7.4) for microscope screening. After the screening, the slides were washed with petroleum ether and xylene (3 min each), dried in an ethanol series (70–80–96%) and subsequently used for FISH (Fig. S6).

*Microscopy and image processing*

The Images of mitotic spreads were captured by an Axio Imager Z2 microscope (Zeiss) with the automatic Metafer-MSearch scanning platform and a CoolCube 1 b/w digital camera (MetaSystems). IKAROS and ISIS imaging programs (Metasystems) were used to analyse both greyscale (Giemsa staining) and colour (post-FISH) images (Fig. S6).

#### Figures

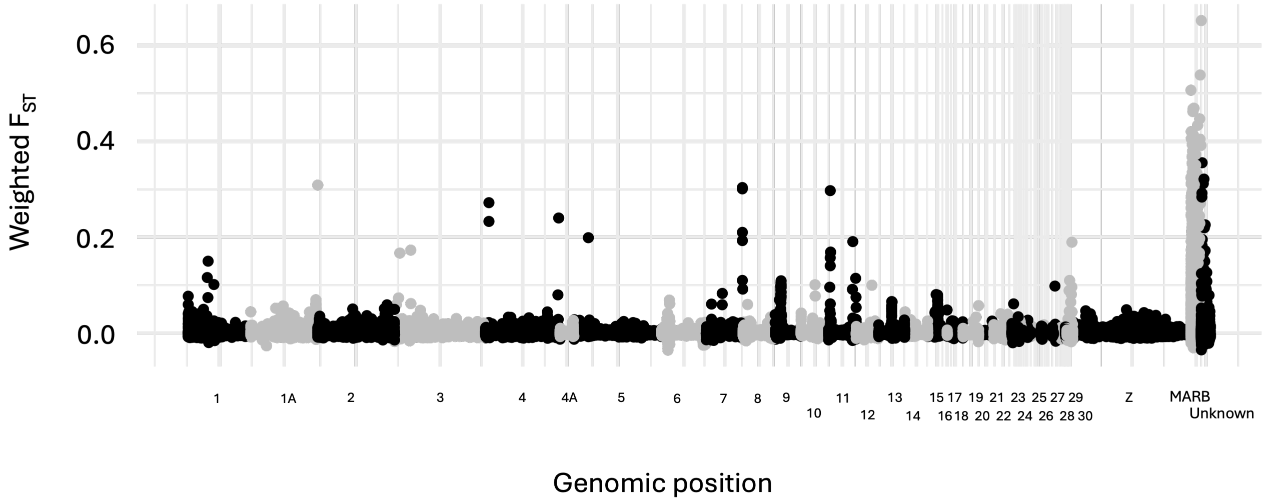

Figure S1. Weighted $F_{ST}$ across the whole genome in 10 kb windows between willow warblers classified as MARB-a present (n = 38) and MARB-a absent (n=38). The classification was made based on the established qPCR assay (1).

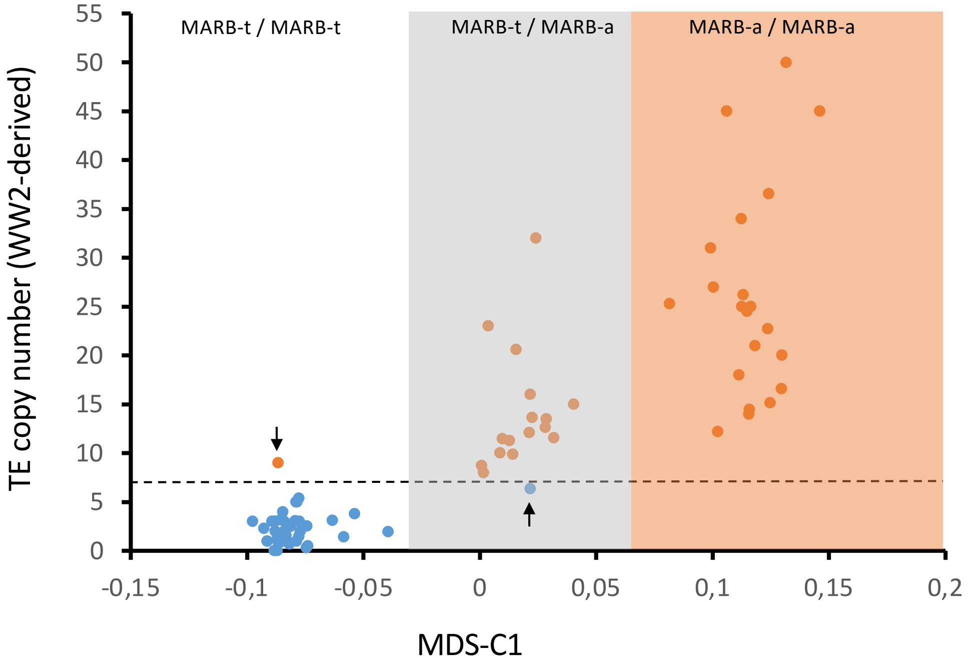

**Figure S2.** MARB-MDS. The relationship between a qPCR-based quantification of TE copy numbers (WW2-derived) and MDS classification of SNPs on MARB-scaffolds (*trochilus* reference genome) for 76 whole-genome re-sequenced willow warblers. The horizontal stippled line shows the selected value separating MARB-a absence (blue dots, <7 copies of WW2-derived TE) from MARB-a presence (orange dots, >7 copies of WW2-derived TE). Arrows point to two individuals who have conflicting classifications with the two methods. The MARB-genotypes have been inferred based on the MDS clusters and are shown with coloured backgrounds.

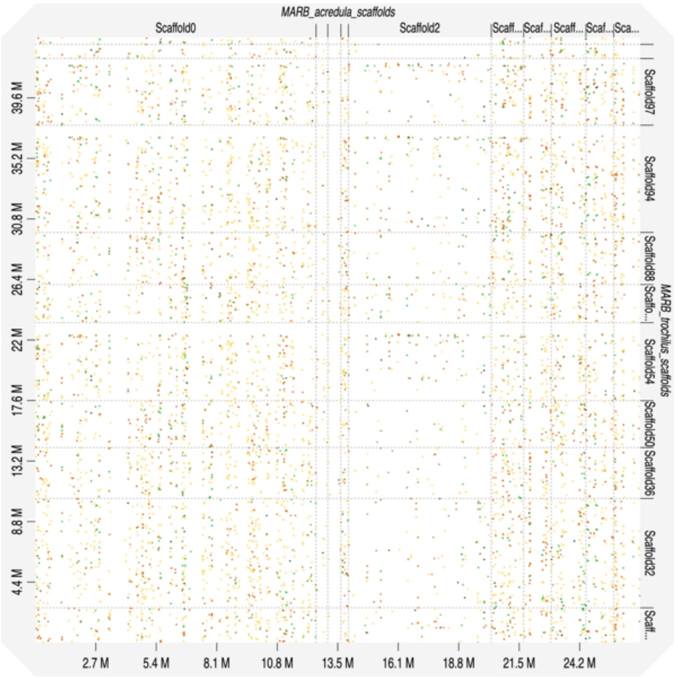

**Figure S3.** Dotplot comparing homology between MARB-a and MARB-t.

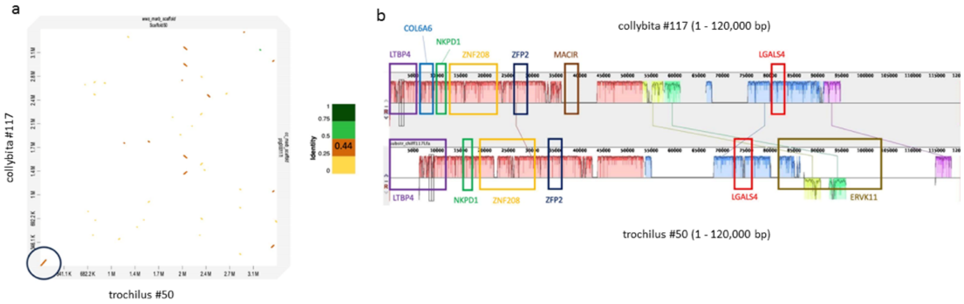

**Figure S4.** Comparison of MARB scaffolds in *trochilus* (#50) and *collybita* (#117). Dotplot of full length (a) and Mauve alignment over the first 120 kb (b). Annotated genes illustrated in coloured boxes.

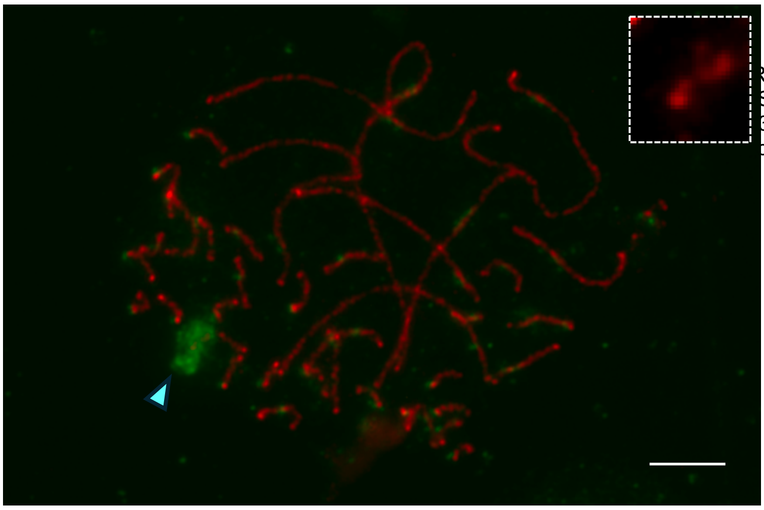

REST antibody in green (blue arrow head). The
SC. The slides were treated with a combination
35, Antibodies Incorporated) antibodies. Anti-
on 1:200, A-11014, Invitrogen). Scale bar = 10

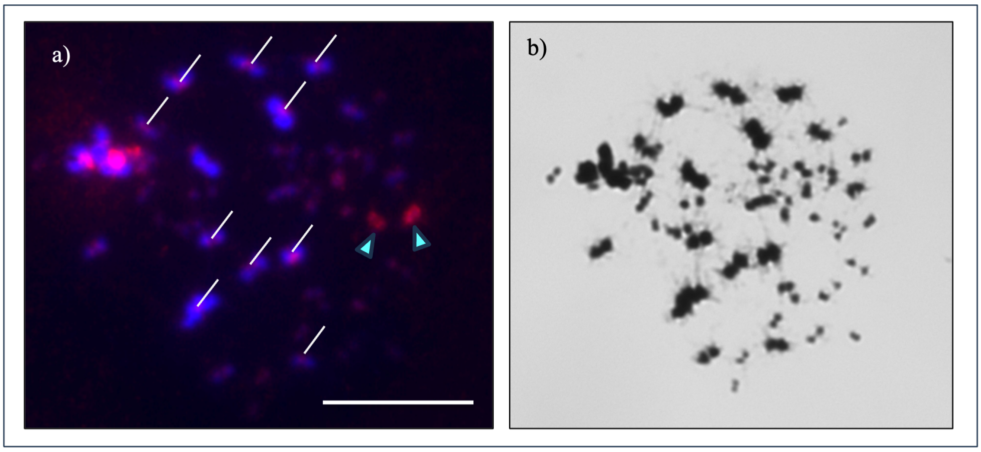

**Figure S6.** Mitotic chromosome spread from willow warbler bone marrow. a) DNA dyed blue with DAPI. In red, probe fragments binding to centromeric and/or pericentromeric regions. The blue arrows point at the MARB chromosome pair, and the white lines point at the most visible centromeric regions. b) Giemsa image of the same spermatocyte. Scale bar = 10 µm.

### Tables

Table S1. Summary statistics for scaffolds identified as MARB in genomes of *acredula*, *trochilus*, *collybita* and *viridanus*. Distance (highlighted blue for MARB, in yellow for the rest of the genome) is the mean bp between the OR's or the repeat elements (rnd-3_fam-21, rnd-3_fam52 and fAlbSat6), respectively. Ratio is the yellow cell divided by the blue cell and is a measure of the density in MARB relative the rest of the genome.

| Taxon | Scaffold | Length (bp) | GC (%) | Unknown (N's) | WW2 derived | OR's | rnd-3_fam-21 | rnd-3_fam-52 | fAlbSat6 |
|---|---|---|---|---|---|---|---|---|---|
| P.t.acredula | Scaffold0 | 12,491,261 | 51.0 | 5,701,254 | 5 | 114 | 495 | 1455 | 862 |
| P.t.acredula | Scaffold2 | 6,354,106 | 51.3 | 2,009,486 | 1 | 43 | 402 | 803 | 423 |
| P.t.acredula | Scaffold5 | 3,990,820 | 51.1 | 1,798,918 | 0 | 39 | 162 | 501 | 286 |
| P.t.acredula | Scaffold21 | 1,853,796 | 51.1 | 601,206 | 0 | 16 | 96 | 302 | 181 |
| P.t.acredula | Scaffold23 | 2,507,465 | 51.1 | 736,825 | 0 | 30 | 137 | 382 | 194 |
| P.t.acredula | Scaffold35 | 1,441,703 | 50.4 | 215,863 | 2 | 17 | 84 | 239 | 135 |
| P.t.acredula | Scaffold55 | 1,227,042 | 50.5 | 621,046 | 1 | 6 | 57 | 116 | 80 |
| P.t.acredula | Scaffold60 | 1,545,618 | 50.7 | 521,912 | 0 | 17 | 66 | 198 | 134 |
| P.t.acredula | Scaffold72 | 1,237,234 | 50.1 | 497,423 | 0 | 9 | 47 | 141 | 86 |
| P.t.acredula | Scaffold81 | 1,173,633 | 51.0 | 642,128 | 0 | 5 | 61 | 107 | 60 |
| P.t.acredula | Scaffold118 | 519,371 | 48.4 | 268,073 | 1 | 2 | 24 | 39 | 20 |
| P.t.acredula | Scaffold196 | 584,621 | 51.1 | 347,013 | 1 | 4 | 16 | 46 | 33 |
| P.t.acredula | Scaffold201 | 402,822 | 50.5 | 87,833 | 0 | 2 | 26 | 81 | 52 |
| P.t.acredula | Scaffold247 | 333,802 | 49.1 | - | 1 | 7 | 18 | 72 | 40 |
| P.t.acredula | TotalMARB | 35,663,294 | 50.4 | 14,048,980 | 12 | 311 | 1691 | 4482 | 2586 |
| P.t.acredula | DistanceMARB | | | | | 69499 | 12782 | 4822 | 8358 |
| P.t.acredula | GenomeWide | 1,159,353,318 | 42.6 | 44,911,456 | 12 | 992 | 4315 | 13189 | 7247 |
| | Non-MARB | 1,123,690,024 | | 30,862,476 | 0 | 681 | 2624 | 8707 | 4661 |
| | DistanceNon-MARB | | | | | 1604739 | 416474 | 125511 | 234462 |
| | Ratio | | | | | 23 | 33 | 26 | 28 |
| P.t.trochilus | Scaffold18 | 2,535,182 | 50.8 | 0 | 0 | 55 | 527 | 167 | 324 |
| P.t.trochilus | Scaffold32 | 7,950,085 | 50.6 | 46 | 0 | 178 | 533 | 1752 | 1014 |
| P.t.trochilus | Scaffold36 | 3,699,632 | 50.5 | 0 | 0 | 73 | 249 | 804 | 447 |
| P.t.trochilus | Scaffold50 | 3,410,838 | 51.2 | 86,750 | 0 | 70 | 224 | 744 | 435 |
| P.t.trochilus | Scaffold54 | 5,674,025 | 51.0 | 0 | 0 | 93 | 294 | 863 | 563 |
| P.t.trochilus | Scaffold57 | 2,774,696 | 50.7 | 38,065 | 0 | 45 | 152 | 542 | 294 |
| P.t.trochilus | Scaffold88 | 3,792,040 | 50.9 | 0 | 0 | 85 | 284 | 806 | 475 |
| P.t.trochilus | Scaffold94 | 7,797,844 | 51.0 | 70,180 | 0 | 140 | 417 | 1368 | 826 |
| P.t.trochilus | Scaffold97 | 4,843,838 | 50.6 | 0 | 0 | 82 | 281 | 841 | 512 |
| P.t.trochilus | Scaffold98 | 1,032,061 | 50.4 | 0 | 0 | 14 | 65 | 182 | 96 |
| P.t.trochilus | Scaffold99 | 511,249 | 49.9 | 0 | 0 | 14 | 33 | 115 | 55 |
| P.t.trochilus | TotalMARB | 44,021,490 | 50.7 | 195,041 | 0 | 849 | 3059 | 8184 | 5041 |
| P.t.trochilus | DistanceMARB | | | | | 51621 | 14327 | 5355 | 8694 |
| P.t.trochilus | GenomeWide | 1,249,322,907 | 43.8 | 2,591,520 | 0 | 1402 | 4765 | 15363 | 8417 |
| | Non-MARB | 1,205,301,417 | | 2,396,479 | | 553 | 1706 | 7179 | 3376 |
| P.t.trochilus | DistanceNon-MARB | | | | | 2175235 | 705103 | 167559 | 356311 |
| | Ratio | | | | | 42 | 49 | 31 | 41 |
| P.c.collybita | Scaffold32 | 6,700,294 | 50.9 | 0 | 1 | 125 | 442 | 1087 | 640 |
| P.c.collybita | Scaffold54 | 8,192,925 | 50.7 | 0 | 0 | 135 | 379 | 1328 | 734 |
| P.c.collybita | Scaffold79 | 6,110,399 | 50.6 | 0 | 1 | 136 | 443 | 1153 | 697 |
| P.c.collybita | Scaffold86 | 2,303,672 | 50.5 | 0 | 2 | 52 | 170 | 430 | 262 |
| P.c.collybita | Scaffold107 | 1,215,101 | 51.2 | 0 | 1 | 24 | 95 | 261 | 159 |
| P.c.collybita | Scaffold117 | 3,460,923 | 52.0 | 0 | 0 | 69 | 255 | 896 | 495 |
| P.c.collybita | TotalMARB | 27,983,314 | | 0 | 5 | 541 | 1784 | 5155 | 2987 |
| P.c.collybita | DistanceMARB | | | | | 51725 | 15686 | 5428 | 9368 |
| P.c.collybita | GenomeWide | 1,247,501,225 | 43.5 | 0 | 5 | 687 | 2893 | 8577 | 4421 |
| | Non-MARB | 1,219,517,911 | | 0 | 0 | 146 | 1109 | 3422 | 1434 |
| P.c.collybita | DistanceNon-MARB | | | | | 8352862 | 1099655 | 356376 | 850431 |
| | Ratio | | | | | 161 | 70 | 66 | 91 |
| P.viridanus | ptg000011 | 6,657,016 | 53.2 | 0 | 0 | 143 | 364 | 1019 | 674 |
| P.viridanus | ptg000023 | 5,725,187 | 52.5 | 0 | 0 | 161 | 402 | 1173 | 751 |
| P.viridanus | ptg000034 | 6,568,268 | 52.6 | 0 | 0 | 172 | 470 | 1236 | 748 |
| P.viridanus | ptg000039 | 12,032,578 | 52.1 | 0 | 0 | 271 | 829 | 1920 | 1222 |
| P.viridanus | ptg000058 | 6,399,663 | 53.2 | 0 | 0 | 102 | 284 | 823 | 521 |
| P.viridanus | ptg000060 | 5,281,243 | 52.9 | 0 | 0 | 115 | 306 | 872 | 496 |
| P.viridanus | ptg000120 | 4,125,993 | 52.2 | 0 | 0 | 108 | 253 | 787 | 512 |
| P.viridanus | TotalMARB | 46,789,948 | | 0 | 0 | 1072 | 2908 | 7830 | 4924 |
| P.viridanus | DistanceMARB | | | | | 43647 | 16090 | 5976 | 9502 |
| P.viridanus | GenomeWide | 1,238,746,615 | 44.0 | 0 | 0 | 1485 | 4634 | 12403 | 7279 |
| | Non-MARB | 1,191,956,667 | | 0 | 0 | 413 | 1726 | 4573 | 2355 |
| P.viridanus | DistanceNon-MARB | | | | | 2886094 | 690589 | 260651 | 506139 |
| | Ratio | | | | | 66 | 43 | 44 | 53 |

**Table S2.** Detailed sequencing procedure of all willow warbler individuals used for the MDS analysis. The read depth measurements correspond to the filtered vcf files.

| Country | Number of individuals | Sequencing | Read depth (mean) | Read depth (Range) |
|---|---|---|---|---|
| Sweden | 32 | NovaSeq s4 flowcell, paired-end of 150 bp read length and v1 sequencing chemistry | 31,59X | 12,87 - 58,46 |
| Sweden | 16 | NovaSeq s4 flowcell, paired-end of 150 bp read length and v1.5 sequencing chemistry | 31,54X | 24,47 - 42,45 |
| Poland | 15 | Novaseq X Plus system, 10B flow cell and XLEAP-SBS sequencing chemistry | 41,01X | 25,02 - 51,98 |
| Lithuania | 13 | Novaseq X Plus system, 10B flow cell and XLEAP-SBS sequencing chemistry | 32,60X | 26,26 - 40,51 |
